# Evidence for a growth zone for deep subsurface microbial clades in near-surface anoxic sediments

**DOI:** 10.1101/2020.03.24.005512

**Authors:** Karen G. Lloyd, Jordan T. Bird, Joy Buongiorno, Emily Deas, Richard Kevorkian, Talor Noordhoek, Jacob Rosalsky, Taylor Roy

**Affiliations:** University of Tennessee, Knoxville, TN USA

## Abstract

Global marine sediments harbor a large and highly diverse microbial biosphere, but the mechanism by which this biosphere is established during sediment burial is largely unknown. During burial in marine sediments, concentrations of easily-metabolized organic compounds and total microbial cell abundance decrease steadily. However, it is unknown whether some microbial clades increase with depth, despite the overall trend of abundance decrease. We show total population increases in 38 microbial families over 3 cm of sediment depth in the upper 7.5 cm of White Oak River (WOR) estuary sediments. Clades that increased with depth were more often anaerobic, uncultured, or common in deep marine sediments relative to those that decreased. Minimum turnover times (which are minimum *in situ* doubling times of growth rates) were estimated to be 2-25 years by combining sedimentation rate with either quantitative PCR (qPCR) or the product of the Fraction Read Abundance of 16S rRNA genes and total Cell counts (FRAxC). Turnover times were within an order of magnitude of each other in two adjacent cores, as well as in two laboratory enrichments of Cape Lookout Bight (CLB), NC, sediments (average difference of 28 ± 19%). qPCR and FRAxC in WOR cores and FRAxC in CLB incubations produced similar turnover times for key deep subsurface uncultured clades *Bathyarchaeota* (8.7 ± 1.9 years) and *Thermoprofundales*/MBG-D (4.1 ± 0.7 years). We conclude that common deep subsurface microbial clades experience a narrow zone of growth in shallow sediments, offering an opportunity for natural selection of traits for long-term subsistence after resuspension events.

**Significance statement:** The current dogma is that the deeply-branching uncultured microbes that dominate global marine sediments do not actually increase in population size as they are buried in marine sediments – rather they exist in a sort of prolonged torpor for thousands of years. This is because no evidence has ever been found that these clades actually increase population sizes, or grow, as they are gradually buried. We discovered that they actually do increase population sizes during burial, but only in the upper few centimeters. This changes our dogma about marine sediments as a vast repository of non-growing microbes, to a vast repository of non-growing microbes with a thin and relatively rapid area of growth in the upper 10 centimeters.

## Introduction

Marine sediments harbor one of the largest microbiomes on Earth, comprising ∼2.9 × 10^29^ microbial cells (1). In most marine sediments, total microbial cells decrease with depth in a log-log relationship since the energy available to support the microbial community decreases as resources are depleted (1, 2). Therefore most microbes in marine sediments are nutrient-limited and in a state of near-zero growth (3, 4). Bulk sedimentary microbial communities have been calculated to have turnover times on the order of tens to hundreds of years, based on microbial respiration rates and energetic requirements (5, 6). The assembly of these deep subsurface communities over many meters of burial appears to occur through selective survival (7) or environmental filtering (8), rather than net growth. Only the most abundant organisms, or those best suited to the changing redox regime, persist while all other clades die off (7–10). However, marine sediment microbes appear to have adaptations to marine sediments, since enzyme affinity (lower K_m_ values) to carbon substrates increases with depth and specialization shifts to highly-degraded substrates with depth (11). These adaptations imply that heritable natural selection has occurred in a sedimentary environment, which implies that these common subsurface clades experience net growth, rather than just persistence, at some point during burial. This net growth may occur in the upper few centimeters of sediments, since microbes have been shown to grow and increase biomass in upper sedimentary layers (12, 13). However, the methods used in these growth studies could not distinguish between different clades nor whether this growth resulted in net population increases with increasing depth. We therefore hypothesized that growth occurs for common subsurface clades in the upper few centimeters of marine sediments, over relatively short time intervals of a few tens of years. This growth would have been missed by previous studies that demonstrated persistence rather than growth of subsurface clades during burial (7– 10) because these studies were conducted in deeper sediments, after net growth had largely ceased.

Microbial populations in marine sediments are phylogenetically diverse (14), so clades may differ in their response to burial. One option is that the cells from each clade die-off at similar rates, with no evidence of growth (Fig. 1). The second option is that some clades experience growth as they are buried, allowing them to increase their population size, even as whole-community cell numbers decrease (Fig. 1). In the second option, the rate of increase in total population size, or turnover time, therefore represents a minimum growth rate estimate. The maximum growth rate could be even higher since growth and death rates may be in equilibrium at each depth layer. Marine sediments contain many phylogenetically divergent non-cultured cells (PDNC) belonging to high-level taxonomic groups with no cultured representatives (15). We hypothesized that 1) some PDNC clades increase in cell abundance with depth in the upper 10 cm of coastal sediments, 2) their turnover times are slower than the < 24 hours doubling times of most microbial cultures, and 3) turnover times are specific to each clade, and are therefore repeatable in different sediment cores and laboratory incubations. We measured cell abundance changes for individual microbial clades with depth in the upper 10 cm (or 40 years of burial time) in the White Oak River (WOR) estuary. Like most marine sediments, they contain diverse PDNCs, and have a steady sediment deposition rate allowing a conversion between depth and time (16). As a second test, we measured the growth of these organisms in 2.2 year-long incubations of sediments from Cape Lookout Bight (CLB) without any nutrient additions so that time could be measured directly (17).

**Figure 1.**
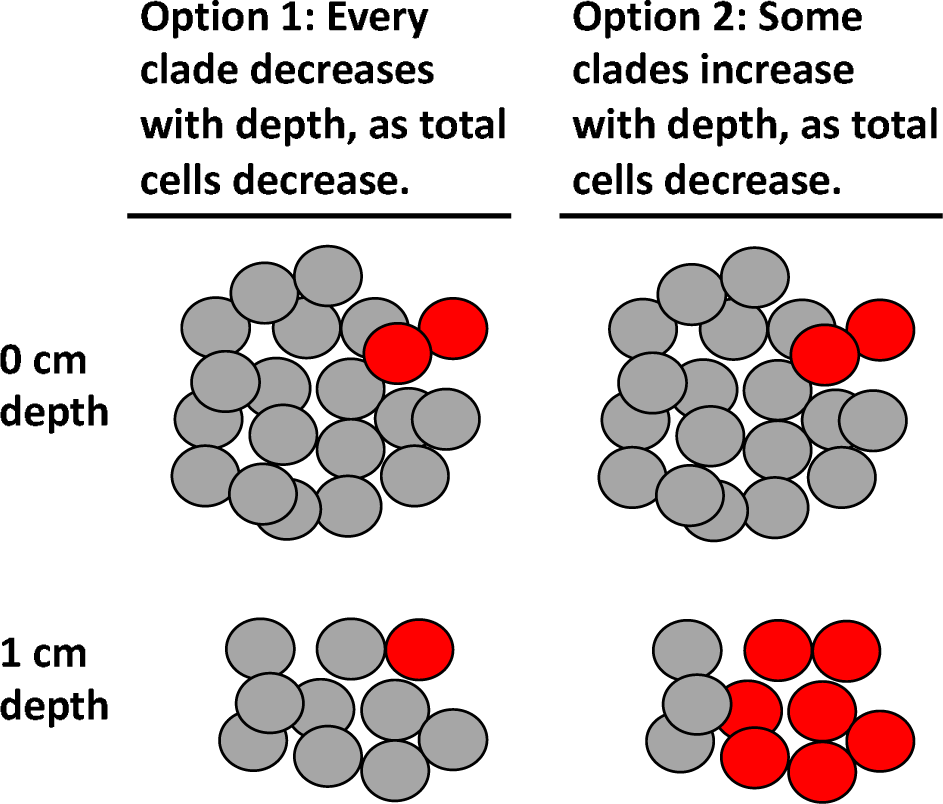
Options for how an individual clade of microbes (red cells) either decrease (option 1) or increase (option 2) with total population decrease with 1 cm sediment depth.

Measuring microbial growth rates in natural marine sediments is not straightforward. Quantification methods, such as catalyzed reporter deposition fluorescent in situ hybridization (CARD-FISH) and quantitative PCR (qPCR) are often inaccurate in marine sediments, due to problems with permeabilizing cells and amplification biases (18, 19). These methods are also limited because they only measure the microbial groups targeted by the chosen probes or primers. Measuring the relative abundance of 16S rRNA gene sequence reads amplified from marine sediments encompasses a wider range of phylogenetic groups, but also does not provide accurate quantifications since DNA extraction methods and primer sequences are biased (20).

However, when considering relative population increases, these biases cancel each other out (Fig. 2). To quantify the relative population changes with depth, one can multiply the <u>F</u>raction of 16S rRNA gene <u>R</u>ead <u>A</u>bundance by the total <u>C</u>ell counts (FRAxC) at each depth or timepoint. The ratio of FRAxC at different depths or timepoints has the bias term in both the numerator and the denominator, so it cancels out, allowing the quantification of relative changes in population size (i.e., a 3-fold increase or a doubling in the example in Fig. 2), but not the exact number of cells that have been gained or lost, over the time interval (18). A second measurement for turnover time was made using qPCR assays, which can also be used to calculate relative population turnover times (18). We found that PDNC and clades that could be inferred to be anaerobic increased with population turnover times of 2-25 years, with good replication for each clade with FRAxC and qPCR, between duplicate cores from the WOR estuary, and between cores from the WOR estuary and CLB sediment incubations. Close relatives of cultures and aerobic organisms decreased with population half lives that were not replicable through the different measurements.

**Figure 2.**
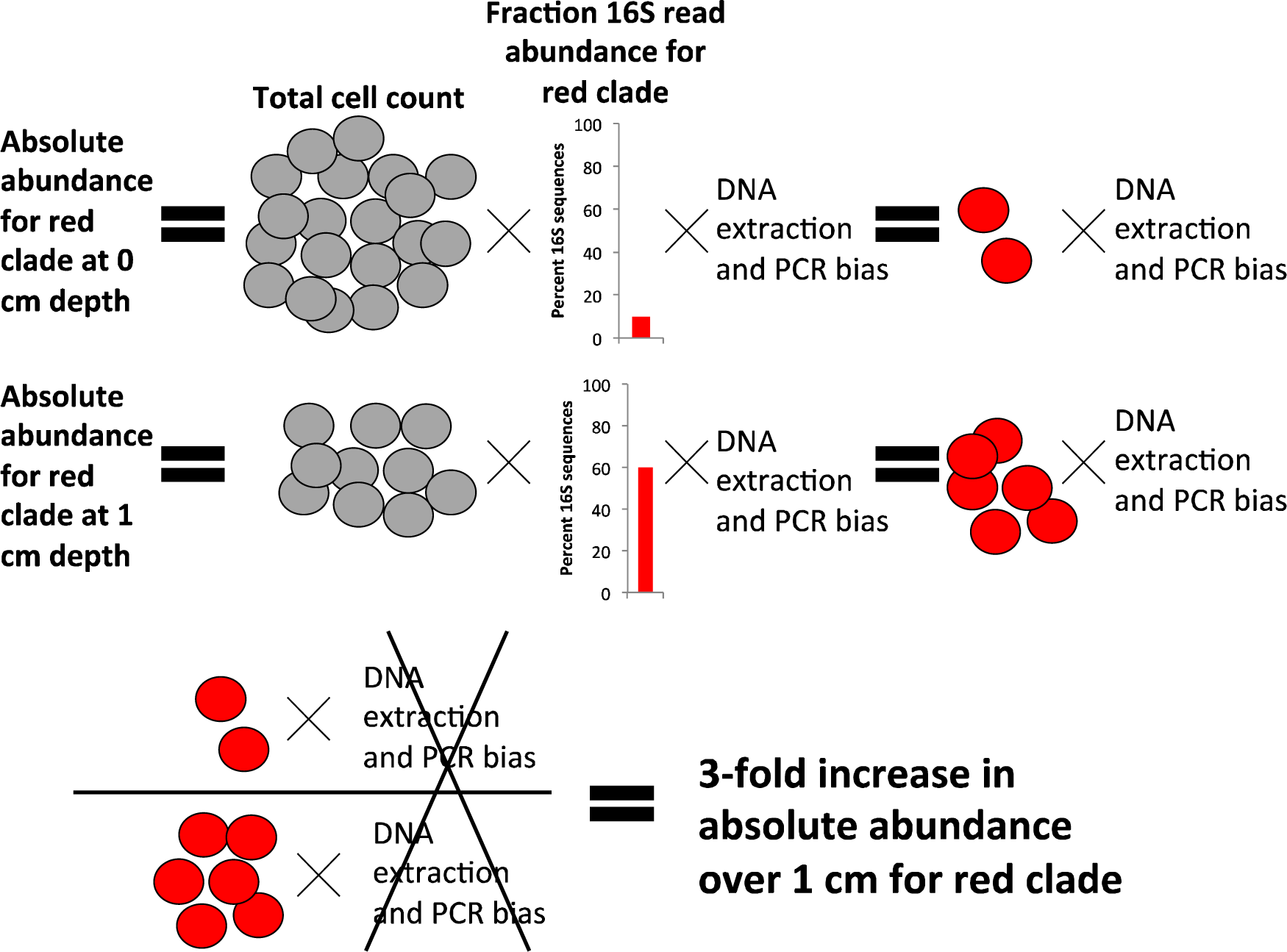
Example to explain how Fraction Read Abundance times Cell abundance (FRAxC) works. The total cell count (grey) decreases with depth. The absolute abundance of an imaginary clade of organisms (red) is equal to the total cell abundance multiplied by the fraction of 16S rRNA gene reads of the red clade and a bias term that cannot be measured accurately and is different for each clade of microbes. Assuming that this bias term is similar over adjacent sediment depths for the same clade of microbes, the relative increase in the red clade with depth can be measured absolutely, since the bias term cancels out.

## Methods

### Sample acquisition and treatment

Four sediment cores of ∼10 cm depth each were retrieved from Station H, WOR estuary (34 44.490’ N, 77 07.44’ W) May 16, 2016, in about 1.5 m water depth, with water salinity of 11 ppt (cores 30, 21, 32, and 34). Each core was sectioned into 1 cm intervals later that day at room temperature (roughly equal to the measured *in situ* water temperature of 25°C) at the University of North Carolina Institute for Marine Sciences. Two cores (cores 30 and 32) were sectioned for qPCR and 16S rRNA gene libraries, two cores (cores 34 and 31, which were taken adjacent to cores 30 and 32, respectively) were sectioned for sulfate concentrations and porosity. The core for cell counts was taken May 28, 2013 (core 7) from the same site (34°44.482’N, 77°7.435’W), and was sectioned in 3 cm intervals. The top 3 cm of marine sediment from CLB (34.6205°N, 76.5500°W) were collected from 10 m water depth on October 2, 2013 and combined into a 2L Erlenmeyer flask that was incubated anoxically for 802 days, as reported previously (17), without any amendment of substrates.

### Geochemistry

To measure sulfate, a 15 ml plastic tube was filled completely with sediment and centrifuged at 5,000x g for 5 minutes. A syringe was used to remove the supernatant below the air interface. The porewater was filtered using a 0.2 μm syringe filter into 100 μl of 10% HCl to a final volume of 1 ml. Porewater sulfate concentrations were determined by ion chromatography (Dionex, Sunnyvale, CA). Porosity was determined by comparing dry and wet weights as described previously (21).

### DNA extraction, qPCR, sequencing, and data analysis

Each 1 cm depth interval of cores 30 and 32 was placed into a Whirlpak bag and frozen immediately on dry ice for later DNA extraction after storage at -80°C. DNA was extracted from frozen sediments (MoBio RNA Powersoil kit with DNA accessory) and enumerated on a NanoDrop 3000. At the timepoints from the CLB incubations, DNA was extracted from frozen samples using the FastDNA kit for Soil (MP Bio, Santa Ana, CA). For both the WOR cores and the CLB incubations, the V4 region of each DNA extract was amplified using the universal primers 515f and 806r (22), prepared via Nexterra kit and sequenced at the Center for Environmental Biotechnology at the University of Tennessee (Knoxville, TN) on an Illumina MiSeq. 16S rRNA gene reads were processed, chimera-checked, and classified via Silva taxonomy (v126; (23)) in mothur (24). All calculations were performed on clades at the fifth taxonomic level, which is roughly the family level, only from clades with >150 total reads in the Miseq run, which left 327 clades from core 30 and 288 clades from core 32. 16S rRNA gene reads were deposited at the Short Read Archive (SRA) at NCBI’s Genbank with BioProject ID PRJNA614649 for WOR sediments and PRJNA321388 for CLB incubations.

Quantitative PCR (qPCR) was used to determine the 16S rRNA gene copy numbers of bacteria, archaea, *Bathyarchaeota*, and Marine Benthic Group – D in the *Thermoplasmata* (with the latter for core 32 only) using the Quantifast SYBRGreen kit (Qiagen) on a BioRad Opticon2 thermocycler. Since absolute copy numbers cannot be accurately measured for DNA in marine sediments (18, 25), only the relative changes in copy number were measured. Our primers were MCG528F and MCG732R for *Bathyarchaeota* (26), MBG-D322F and MBG-D569R for Marine Benthic Group – D (27), ARCH915F (28) and ARC1059R (29) for archaea, and BAC340F and BAC515R for bacteria (30).

### Population turnover time calculations

Population turnover times for a family-level classification were calculated from relative 16S rRNA gene abundances and total cell counts as described previously (17), with the following relationship:

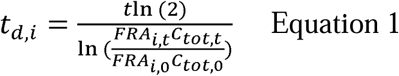

where *t*_*d,i*_ is the doubling time of the *i*th family clade, *t* is the elapsed time, *FRA*_*i,t*_ is the fraction read abundance of the *i*th clade at time *t*, and *C*_*tot,t*_ is the total number of cells at time *t*. The product of the fraction 16S rRNA gene abundance of a particular clade and the total cell abundance will be referred to as the FRAxC of a particular clade. To calculate the population turnover for a specific group of target organisms using qPCR, we used the following relationship:

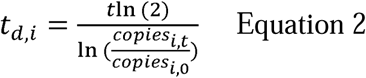

where *copies* is the number of copies of target 16S rRNA gene quantified via qPCR. Equations 1 and 2 are equivalent to ln(2) divided by the slope time vs. the natural log of *FRA*_*i*_*C*_*tot*_ or the natural log of *copies*_*i*_. For incubations from CLB, time from the start of the experiment was known. For samples taken from the WOR estuary, time at each depth was determined based on an age model (31), using the porosity measurements for core 31 and a sedimentation rate of 0.26 cm/year (32).

### Cell counts

Sediments were fixed in 3% paraformaldehyde for a few hours, washed twice with PBS, and stored at -20°C in PBS:ethanol. Sediments were sonicated at 20% power for 40 seconds, diluted to 1:40 in PBS, filtered onto a 0.2 μm polycarbonate filter (Fisher Scientific, Waltham, MA), stained with SYBR Gold (Invitrogen, Carlsbad, CA), and counted at 100x magnification on an epifluorescent microscope. To interpolate cell counts into 1 cm intervals, an exponential curve fit was applied to the samples within the 0-10 cm interval with the following fit: depth = 137.77 exp(−1×10^−9^ x cells/ml), R^2^=0.94. Two more cell count curves were used to test the sensitivity of turnover times to cell count profile variations. In one, cell counts from a 2012 WOR core were previously published (17), and resulted in the following curve fit over 0-10 cm: depth = 14.13 exp(−4×10^−10^ x cells/ml), R^2^=0.93. In the second, cell counts were taken using WebPlotDigitizer from previously published estuarine sediments from Ashleworth Quay (33). For the CLB incubations, SYBR Gold direct counts of paraformaldehyde-fixed cells were performed as described previously (17).

## Results and Discussion

Sulfate concentrations and porosity were constant with depth in the upper 5 cm of core 34 and the upper 3 cm of core 31 (Fig. S1) due to bioirrigation and bioturbation. Below this, sulfate concentrations decreased due to microbial sulfate reduction as has been observed in other cores at this site (21, 34). Total cell abundance decreased with depth (Fig. S2). The FRAxC and qPCR of total archaea, *Bathyarchaeota* (previously called MCG or MG 1.3 (35)), and *Thermoprofundales* (previously called MBG-D (36)) increased slightly below the depth of bioirrigation in both WOR cores (Fig. S3). The FRAxC and qPCR of total bacteria decreased with depth in both cores. qPCR values were lower than FRAxC values, demonstrating that these methods are subject to amplification biases and are best used to measure relative changes rather than absolute values (18). The turnover times measured with FRAxC and qPCR methods were within a year of each other for *Bathyarchaeota* and MBG-D (Table 1), suggesting that FRAxC is as good a measure of relative abundance as qPCR. FRAxC, however, opens the possibility of measuring turnover times for any clade represented in the 16S rRNA gene libraries, without developing primer sets for each one.

**Table 1.** Doubling times for all clades that increased in both cores of White Oak River (WOR) estuary sediments and replicate incubations from the Cape Lookout Bight (CLB) marine sediments. Data are expressed in years. FRAxC denotes values calculated by the increase of the product of 16S rRNA gene read percentages and total cells over depth (WOR) or time (CLB). qPCR denotes values calculated by the increase of qPCR of 16S rRNA genes over depth. Hyphen indicates no data because qPCR was not attempted for that clade or sample.

| Clade | WOR marine sediment |  |  |  | CLB marine sediment |  |
| --- | --- | --- | --- | --- | --- | --- |
|  | <u>Core 30</u> |  | <u>Core 32</u> |  | <u>Incubation 2</u> | <u>Incubation 3</u> |
|  | FRAxC | qPCR | FRAxC | qPCR | FRAxC | FRAxC |
| *Bathyarchaeota*, or MCG | 10.1 | 10.3 | 8.9 | 10.0 | 6.4 | 6.3 |
| Group C3, MCG-related | 16.2 | - | 14.9 | - | 2.1 | 1.8 |
| *Thermoprofundales* (MBG-D) | 4.6 | - | 3.4 | 4.8 | 3.4 | 4.4 |
| *Thermoprofundales* (MG-III) | 3.7 | - | 2.8 | - | 1.4 | 1.5 |
| *Thermoprofundales* (20a-9) | 7.1 | - | 3.7 | - | 0.8 | 0.7 |

Using FRAxC in the WOR cores, the doubling time (defined as a positive value for turnover time) or half-life (defined as negative a value for turnover time) was calculated for each family-level clade across the three centimeters below the bioirrigated upper layer (5.5-7.5 cm for core 30 and 3.5-5.5 cm for core 32, Table S1). These three depths produced the highest number of clades that increased with depth. A similar increase in the relative contribution from anaerobic organisms was observed immediately below the bioirrigation zone in the marine sediments of Aarhus Bay, Denmark (8). Of the 315 clades, 278 were present in both cores, and 60% of these had R^2^ > 0.4 to the curve fit to equation 1, meaning that a change in FRAxC over depth could be observed. The lenient cut-off of R^2^ > 0.4 was chosen in order to perform downstream analyses on the largest possible selection of clades. Of these, 80% had doubling times or half-lives between 2 and 25 years. This means that the sampling interval of ∼8 years was sufficient to resolve clades that underwent at least a third of one doubling or halving in that interval. Clades that increased faster or more slowly with depth were outside the detection limit. Read abundance between different clades did not correlate with turnover time (Fig. S4). In other words, a clade’s relative abundance in the 16S rRNA gene libraries did not predict whether it would increase or decrease with depth.

To determine how sensitive turnover times were to differences in the cell abundance curves used to calculate FRAxC, comparisons were made with cell counts from a 2012 WOR core (17) and estuarine sediments from Ashleworth Quay (33). For core 30, 92% and 100% of the clades with turnover times of 2-25 years still fell within that range using the 2012 and Ashleworth Quay cell counts. For core 32, these numbers were 98% and 100%. This suggests that this method is fairly robust to variations in cell abundance curves.

The direction of either increase or decrease was replicable in WOR cores 30 and 32 for 131 out of 133 clades (Fig. 3a). Clades that increased with depth were common marine subsurface inhabitants: *Chloroflexi, Deltaproteobacteria, Planctomycetes, Bathyarchaeota, Lokiarchaeota*, MBG-D, etc (37). Those that decreased with depth were common seawater inhabitants: *Betaproteobacteria, Alphaproteobacteria, Gammaproteobacteria, Thaumarchaeota* (MG-I), *Bacteroidetes, Acidobacteria* (38). Clades from uncultured phyla were more likely to increase rather than decrease with depth (9/11 clades). Clades from cultured orders, families, genera, or species were more likely to decrease rather than increase with depth (57/60 clades). This suggests that easily cultured organisms were dying off and difficult-to-culture clades, or PDNC, were growing. Aerobes and nitrate reducers were more likely to decrease with depth (23/23 clades where all cultures are obligate aerobes or nitrate reducers, Fig. 3a, inset). Doubling times for clades that increased were well-correlated for individual clades between the two WOR cores (slope = 0.87, R^2^ = 0.64), suggesting that they may have been growing at rates determined by their particular physiological traits. The mean deviation from the mean between the duplicate cores was 28 ± 19%. This means that turnover times were accurate at least within an order of magnitude. For instance, a clade with a turnover time of 10 is very likely to have an actual turnover time within at least a range of 5-15 years. In contrast, half-lives for clades that decreased were poorly correlated for individual clades between cores (slope = 0.27, R^2^ = 0.11). Clades that decreased with depth may have been dying or decaying because they could not meet their energetic requirements in this anoxic environment, and, unlike the turnover times of the growing organisms, their rates of decay/death were not specific to their physiologies. These results support the conclusion that FRAxC doubling times represent real population changes during burial and not random variations.

**Figure 3.**
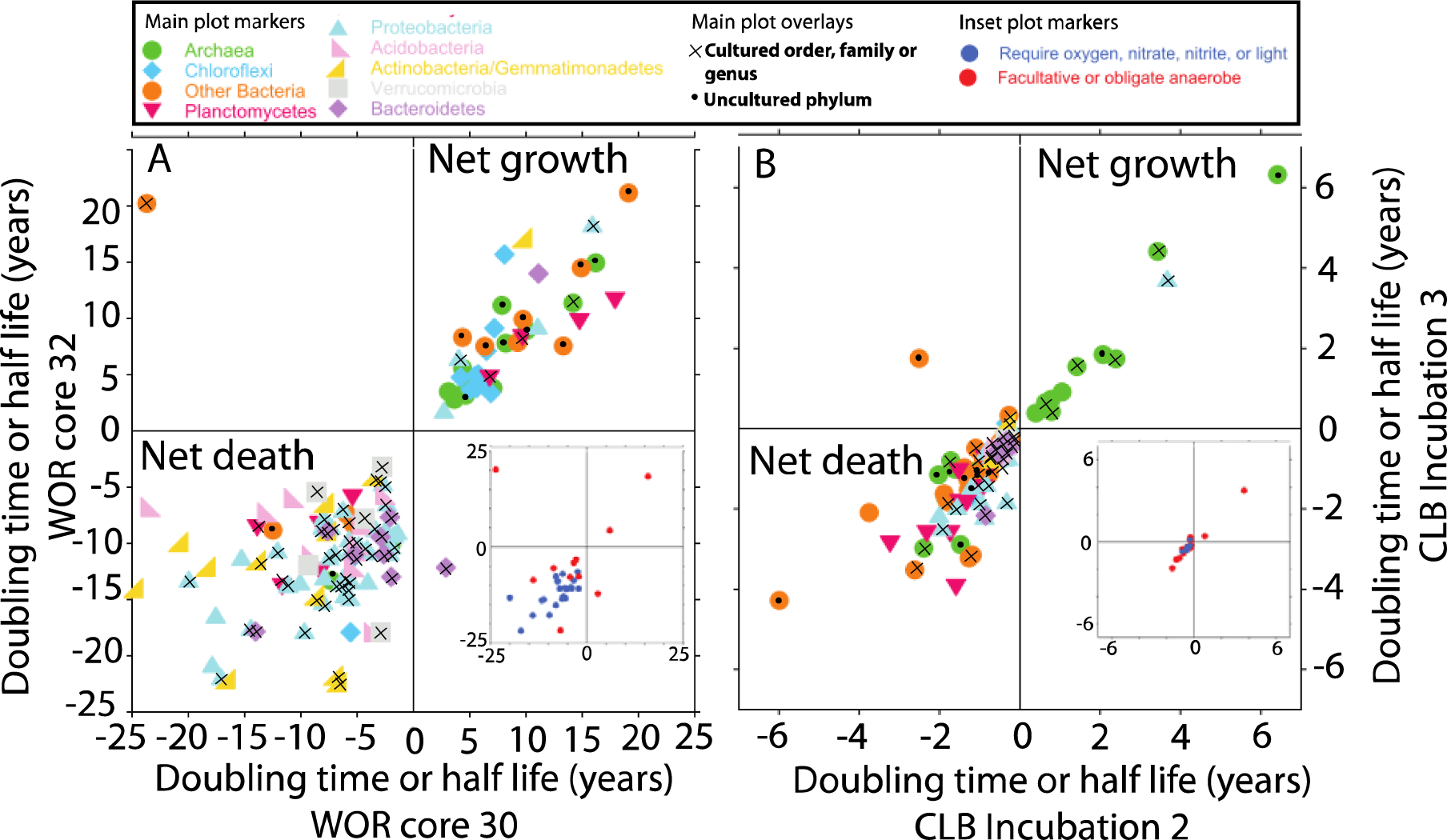
Multi-year doubling times for uncultured microbes in marine sediments, measured in A) down-core trends in White Oak River estuary or B) incubation time course experiments from Cape Lookout Bight. In each panel, replicates are plotted against each other to show good correlation between clades that increase (upper right quadrants) or decrease (lower left quadrants) with depth. In A) clades that grow tend to be common subsurface clades (in dark colors) and uncultured phyla (black dots); clades that decay tend to be common seawater clades (in pastels) and more closely related to cultures (x’s). Insets show the subset of clades belonging to cultured families where all cultured relatives are oxygen- or nitrate-reducing (blue), or contain at least some non-nitrogen-cycling facultative anaerobes (red). Note different timescales for WOR and CLB data.

As a separate check on the feasibility of these turnover times, we used the depth-integrated sulfate reduction rate (0.93 µM cm^-3^ d^-1^, Fig. S1) to estimate a turnover time for sulfate reducers of 12 years, assuming that 10% of total cells were sulfate reducers with growth yields of 2 g cell C/mol (3), and cell carbon contents of 23 fg/cell (39). This falls well within our results from FRAxC of 2-25 years doubling times, including one clade with cultured sulfate reducers, *Desulfarculaceae*, that had doubling times of 15 and 18 years in the two cores. Previous work in arctic and temperate marine sediments also supports generation times of over a year in marine sediment incubations where total cell abundance of sulfate reducing bacteria held constant despite increasing metabolic rates over a year-long incubation (40).

Should the population turnover times estimated here approximate the growth rate of these clades, then these microbes would be considered dormant relative to pure cultures (4). However, if the populations are in steady state with some resource that gradually increases with depth (such as a lessening influence of bioturbation, for example), then cell replication is roughly balanced by cell death, so an increase in total population size does not necessarily reflect the growth rate, which may be much faster. To estimate how much the growth rate is likely to differ from doubling times, imagine a clade which increased in relative population from 4% to 50% of the population during a drop in total community size of 1,000,000 cells to 100,000 cells over 2 years. This clade would have a doubling time of 6.21 years. During those two years, the net gain was therefore 10,000 cells. If no cells died, then 10,000 divisions occurred and the doubling time is a growth rate. Death rates can vary greatly among different microbial groups (41), but if we estimate the death rate for a clade to be equal to the total community death ratio of 10:1 for this example, then 100,000 divisions would have occurred and the true growth rate would have been 1.16 years. So, it is possible that the actual growth rates of the clades in our study were faster than 2-25 years, but they may not have been orders of magnitude faster if the decay rates are slow for cells adapted to low energy environments.

Two alternative possibilities could also explain an increase with depth in sediment cores, without indicating population growth. First, this increase could represent a mixing curve between a deep source of microbial cells diffusing up toward the surface sediments. Second, the increase could be the result of depositional changes, with greater loading of these subsurface clades in previous depositional events. To rule out these alternative possibilities, we tracked community succession in marine sediment from Cape Lookout Bight, NC, in a laboratory mesocosm under methanogenic conditions over 2.2 years (Fig. 3b). Doubling times or half-lives were calculated with FRAxC for each family level clade using timepoints between 40 and 802 days (40, 47, 54, 61, 75, 80, 86, 94, 107, 114, 122, and 802 days, Table S2). All 601 taxa were present in both incubations. R-squared was not a good method of quality control for the CLB incubation data because the data were not evenly distributed across the time interval. Instead, an abundance cutoff was imposed such that each clade must have had at least one timepoint with a value of at least 14 for ln(FRAxC), which left 195 taxa remaining. Of these, 96%, or 188, had doubling times or half-lives between 0.1 and 7 years (Fig. 3b, Table S2). As with the WOR cores, the sampling interval of 0.1 to 2.2 years was sufficient to resolve clades that underwent a least a third of one doubling or halving in that interval, with slower or faster clades outside the detection limit. Also, similarly to the WOR cores, the direction of increase or decrease was replicable in duplicate incubations (182/188 clades), clades that increased were more similar between replicate incubations (slope = 1.05, R^2^ = 0.96), and clades that decreased were less similar between replicate cores (slope = 0.83, R^2^ = 0.53).

The clades that increased over time in the CLB incubations were largely methane-cycling archaea (*Methanosarcinales, Methanomicrobiales*, and ANME-1), which was expected since the onset of methanogenesis occurred during the sampling interval. The turnover times of methanogen-like archaea of 0.3 to 0.8 years, were consistent with previously measured doubling times of one of these clades, ANME-1, grown in an enrichment, at 0.6 years (42). Of the five clades that increased in both the WOR sediments and the CLB incubations (*Bathyarchaeota*, C3, MBG-D, MG-III, and 20a-9), four increased faster in the CLB incubations, but most had multi-year turnover times in both types of experiments and measurement methods (Table 1). Two clades (*Bathyarchaeota* and MBG-D) had both types of measurements (qPCR and FRAxC) in both types of experiments (WOR cores and CLB incubations). The mean of all these measurements resulted in turnover times of 8.7 ± 1.9 years for *Bathyarchaeota* and 4.1 ± 0.7 years for MBG-d; the relatively small standard deviation suggests that these turnover times are accurate within a few years. Many of the clades with slower turnover times in the WOR cores were present in the CLB incubations, but did not have turnover times that met our quality cut-off. We may have been able to resolve their growth rates if we had waited ∼8 years. The lab incubation experiment therefore also supports a multi-year population turnover time for anoxic uncultured marine subsurface organisms, in an experimental system that rules out diffusional mixing and depositional changes as drivers for the population changes.

These results suggest that the log-log decay rate with depth for the total microbial community does not apply to some clades within the marine sediment biosphere. Some clades experience periods of net growth, resulting in population increases during burial (Option 2 in Fig. 1). This has important implications for microbial adaptations to the marine subseafloor biosphere. Many of these clades have been shown to be dormant or non-growing for most of the sediment column (3, 9). In other environments, such as agricultural soils, periods of dormancy are interspersed with periods of growth (4). Our results suggest that the period of growth for marine sediment microbes may occur in the upper few cm, over a time period that is much shorter than the full marine sediment column, which can be millions of years old.

This suggests a possible mechanism for adaptation to long-term dormancy in marine sediments, building on the Growth Advantage in Stationary Phase (GASP), where *E. coli* cells that have been starved for years outcompete fresh cells under low substrate conditions (43). Our results suggest that these GASP-like adaptations to long-term starvation could also be selected for after resuspension events, allowing GASP to be extrapolated over much longer timescales in marine sediment, as was predicted previously (44). If deep sedimentary cells with subsistence-promoting or GASP-like mutations are resuspended either by a storm or sediment slumping, these organisms would have a growth advantage over others as they were buried a second time through the relatively brief and shallow growth zone. This means that although evolutionary changes do not occur during the process of burial and near-zero cell growth over thousands of years (7), the ability to subsist over thousands of years in marine sediment is a trait that could have been selected for within the growth zone in shallow sediments. Microbes might even be ejected from deeper sediments depths by mud volcanoes into overlying seawater, where they can be carried by currents and re-deposited at the seafloor (45). Our results suggest that such organisms may revive and begin growing shortly after deposition.

If these turnover times are similar to growth rates, then these organisms grow much more slowly than most laboratory cultures, which double in less than a day. This agrees with previous work showing that microbes grow much more slowly in the environment than in laboratory culture. *Staphylococcus aureus* grows 48-fold faster in culture than it does in an infected lung (46) and *Leucothrix mucor* grows 7-fold faster in culture than it does in seawater (47). The fact that so many of the clades that grew in our study belong to uncultured clades suggests that slow growth rates may be a reason why so many of them remain uncultured. Recent successes with culturing slow-growing strains from previously-uncultured phyla *Lokiarchaeota* and *Atribacteria* attest to the necessity of extremely long wait times for growth; 2000 days in one case (48, 49).

Even if it is possible to speed up such organisms sufficiently to culture them, their traits expressed under such conditions are likely to greatly diverge from those expressed in their natural environment. Therefore, even when cultures are available, it is crucial to study these organisms in their natural environmental settings. Using label incorporation (46, 50, 51), or tracking slow growth in a mixed population under different conditions, will allow for growth experiments on the environmentally abundant, evolutionarily distinct, uncultured microbes from resource-limited environments. Such experiments are necessary to assess modifications to cell cycle regulation, proteins, or plasma membranes that enable such extraordinarily slow growth, as well as determine how this narrow growth zone in surficial sediments may select for the traits that enable long-term subsistence of the deep subsurface biosphere.

## Supporting information

All supplemental information

## Acknowledgments

The authors thank Michael Piehler for lab space at the University of North Carolina Institute for Marine Sciences, Frank Löffler, Steve Wilhelm and Erik Zinser for sharing laboratory equipment, Dan Williams, and Ameen Abdel-Khalek for help in lab and with sequencing, and Andreas Schramm for planting the seed of the idea of growth in the upper 10 cm. This material is based upon work supported by (1) the National Science Foundation under grant numbers OCE-1431598 (KL), (2) the NSF Center for Dark Energy Biosphere Investigations (OCE-0939564) contribution # *to be filled in* (KL), (3) the Alfred P. Sloan Foundation Fellowship (FG-2015-65399, KL), (4) the Simons Foundation (404586, KL).

## References

1. Kallmeyer J, Pockalny R, Adhikari RR, Smith DC, D’Hondt S. 2012. Global distribution of microbial abundance and biomass in subseafloor sediment. Proc Natl Acad Sci U S A 109:16213–16216.

2. Parkes RJ, Cragg BA, Wellsbury P. 2000. Recent studies on bacterial populations and processes in subseafloor sediments□: A review. Hydrogeol J 11–28.

3. Hoehler TM, Jørgensen BB. 2013. Microbial life under extreme energy limitation. Nat Rev Microbiol 11:83–94.

4. Lennon JT, Jones SE. 2011. Microbial seed banks: the ecological and evolutionary implications of dormancy. Nat Rev Microbiol 9:119–30.

5. Braun S, Mhatre SS, Jaussi M, Røy H, Kjeldsen KU, Seidenkrantz M, Jørgensen BB, Lomstein BA. 2017. Microbial turnover times in the deep seabed studied by amino acid racemization modelling. Sci Rep 7:5680.

6. Jorgensen BB. 2011. Deep subseafloor microbial cells on physiological standby. Proc Natl Acad Sci 108:18193–18194.

7. Starnawski P, Bataillon T, Ettema TJG, Jochum LM, Schreiber L, Chen X, Lever MA, Polz MF, Jørgensen BB, Schramm A, Kjeldsen KU. 2017. Microbial community assembly and evolution in subseafloor sediment. Proc Natl Acad Sci 114:2940–2945.

8. Marshall IPG, Ren G, Jaussi M, Lomstein BA, Jørgensen BB, Røy H, Kjeldsen KU. 2019. Environmental filtering determines family-level structure of sulfate-reducing microbial communities in subsurface marine sediments. ISME J 1920–1932.

9. Kirkpatrick JB, Walsh EA, D’Hondt S. 2016. Fossil DNA persistence and decay in marine sediment over hundred-thousand-year to million-year time scales. Geology 44:615–618.

10. Jochum LM, Chen X, Lever MA, Loy A, Jørgensen BB, Schramm A, Kjeldsen KU. 2017. Depth Distribution and Assembly of Sulfate-Reducing Microbial Communities in Marine Sediments of Aarhus Bay. Appl Environ Microbiol 83:1–15.

11. Steen AD, Kevorkian RT, Bird JT, Dombrowski N, Baker BJ, Hagen SM, Mulligan KH, Schmidt JM, Webber AT, Royalty T, Alperin MJ. 2019. Kinetics and identities of extracellular peptidases in subsurface sediments of the White Oak River Estuary, NC. Appl Environ Microbiol 85:1–14.

12. Karl D, Novitsky J. 1988. Dynamics of microbial growth in surface layers of a coastal marine sediment ecosystem. Mar Ecol Prog Ser 50:169–176.

13. Novitsky J, Karl D. 1986. Characterization of microbial activity in the surface layers of a coastal subtropical sediment. Mar Ecol Prog Ser 28:49–55.

14. Parkes RJ, Webster G, Cragg BA, Weightman AJ, Newberry CJ, Ferdelman TG, Kallmeyer J, Jørgensen BB, Aiello IW, Fry JC. 2005. Deep sub-seafloor prokaryotes stimulated at interfaes over geological time. Nature 436:390–394.

15. Lloyd KG, Steen AD, Ladau J, Yin J, Crosby L. 2018. Phylogenetically novel uncultured microbial cells dominate Earth microbiomes. mSystems 3:e00055–18.

16. Kubo K, Lloyd KG, F Biddle J, Amann R, Teske A, Knittel K. 2012. Archaea of the Miscellaneous Crenarchaeotal Group are abundant, diverse and widespread in marine sediments. ISME J 6:1949–65.

17. Kevorkian R, Bird JT, Shumaker A, Lloyd KG. 2018. Estimating population turnover rates by relative quantification methods reveals microbial dynamics in marine sediment. Appl Environ Microbiol 84:1–16.

18. Lloyd KG, May MK, Kevorkian RT, Steen AD. 2013. Meta-analysis of quantification methods shows that archaea and bacteria have similar abundances in the subseafloor. Appl Environ Microbiol 79:7790–7799.

19. Buongiorno J, Turner S, Webster G, Asai M, Shumaker AK, Roy T, Weightman A, Schippers A, Lloyd KG. 2017. Inter-laboratory quantification of Bacteria and Archaea in deeply buried sediments of the Baltic Sea (IODP Expedition 347). FEMS Microbiol Ecol fix007.

20. Polz MF, Cavanaugh CM. 1998. Bias in template-to-product ratios in multitemplate PCR. Appl Environ Microbiol 64:3724–3730.

21. Lloyd KG, Alperin MJ, Teske A. 2011. Environmental evidence for net methane production and oxidation in putative ANaerobic MEthanotrophic (ANME) archaea. Environ Microbiol 13:2548–2564.

22. Caporaso JG, Lauber CL, Walters WA, Berg-Lyons D, Huntley J, Fierer N, Owens SM, Betley J, Fraser L, Bauer M, Gormley N, Gilbert JA, Smith G, Knight R. 2012. Ultra-high-throughput microbial community analysis on the Illumina HiSeq and MiSeq platforms. ISME J 6:1621–1624.

23. Pruesse E, Quast C, Knittel K, Fuchs BM, Ludwig W, Peplies J, Glöckner FO. 2007. SILVA: a comprehensive online resource for quality checked and aligned ribosomal RNA sequence data compatible with ARB. Nucleic Acids Res 35:7188–7196.

24. Schloss PD, Westcott SL, Ryabin T, Hall JR, Hartmann M, Hollister EB, Lesniewski RA, Oakley BB, Parks DH, Robinson CJ, Sahl JW, Stres B, Thallinger GG, Horn DJ Van, Weber CF. 2009. Introducing mothur: Open-source, platform-independent, community-supported software for describing and comparing microbial communities. Appl Environ Microbiol 75:7537–7541.

25. Buongiorno J, Turner S, Webster G, Asai M, Shumaker AK, Roy T, Weightman A, Schippers A, Lloyd KG. 2017. Interlaboratory quantification of Bacteria and Archaea in deeply buried sediments of the Baltic Sea (IODP Expedition 347). FEMS Microbiol Ecol 93.

26. Kubo K, Lloyd KG, F Biddle J, Amann R, Teske A, Knittel K. 2012. Archaea of the Miscellaneous Crenarchaeotal Group are abundant, diverse and widespread in marine sediments. ISME J 6.

27. Lloyd KG, Schreiber L, Petersen DG, Kjeldsen KU, Lever MA, Steen AD, Stepanauskas R, Richter M, Kleindienst S, Lenk S, Schramm A, Jørgensen BB. 2013. Predominant archaea in marine sediments degrade detrital proteins. Nature 496:215–218.

28. Stahl DA, Amann R. 1991. Development and application of nucleic acid probes. John Wiley & Sons, Chichester, UK.

29. Yu Y, Lee C, Kim J, Hwang S. 2005. Group-specific primer and probe sets to detect methanogenic communities using quantitative real-time polymerase chain reaction. Biotechnol Bioeng 89:670–9.

30. Nadkarni MA, Martin FE, Jacques NA, Hunter N. 2002. Determination of bacterial load by real-time PCR using a broad-range (universal) probe and primers set. Microbiology 148:257–266.

31. Larowe DE, Amend JP. 2015. Power limits for microbial life. Front Microbiol 6:1–11.

32. Benninger LK, Martens CS. 1983. Sources and fates of sedimentary organic matter in the White Oak and Neuse estuariesWater Resources Research Insitute of the University of North Carolina, UNC-WRRI-83-194.

33. Wellsbury P, Herbert RA, Parkes RJ. 1996. Bacterial activity and production in near-surface estuarine and freshwater sediments 19.

34. Martens CS, Albert DB, Alperin MJ. 1998. Biogeochemical processes controlling methane in gassy coastal sediments — Part 1. A model coupling organic matter flux to gas production, oxid1. Martens CS, Albert DB, Alperin MJ (1998) Biogeochemical processes controlling methane in gassy coastal sedim. Methods 18.

35. Meng J, Xu J, Qin D, He Y, Xiao X, Wang F. 2013. Genetic and functional properties of uncultivated MCG archaea assessed by metagenome and gene expression analyses. ISME J 1–10.

36. Zhou Z, Liu Y, Lloyd KG, Pan J, Yang Y, Gu J-D, Li M. 2018. Genomic and transcriptomic insights into the ecology and metabolism of benthic archaeal cosmopolitan, Thermoprofundales (MBG-D archaea). ISME J.

37. Teske A, Sørensen KB. 2008. Uncultured archaea in deep marine subsurface sediments: have we caught them all? ISME J 2:3–18.

38. Galand PE, Casamayor EO, Kirchman DL, Lovejoy C. 2009. Ecology of the rare microbial biosphere of the Arctic Ocean. Proc Natl Acad Sci U S A 106:22427–32.

39. Braun S, Morono Y, Littmann S, Kuypers M, Aslan H, Dong M, Jorgensen BB, Lomstein BA. 2016. Size and carbon content of sub-seafloor microbial cells at Landsort Deep, Baltic Sea. Front Microbiol 7:1–13.

40. Robador A, Brüchert V, Jørgensen BB. 2009. The impact of temperature change on the activity and community composition of sulfate-reducing bacteria in arctic versus temperate marine sediments. Environ Microbiol 11:1692–1703.

41. Hao X, Wang Q, Zhang X, Cao Y, Mark Loosdrecht CM van. 2009. Experimental evaluation of decrease in bacterial activity due to cell death and activity decay in activated sludge. Water Res 43:3604–3612.

42. Nauhaus K, Albrecht M, Elvert M, Boetius A, Widdel F. 2007. In vitro cell growth of marine archaeal-bacterial consortia during anaerobic oxidation of methane with sulfate. Environ Microbiol 9:187–196.

43. Finkel SE. 2006. Long-term survival during stationary phase: evolution and the GASP phenotype. Nat Rev Microbiol 4:113–20.

44. Lever MA, Rogers KL, Lloyd KG, Schink B, Thauer RK, Hoehler TM, Jørgensen BB. 2015. Life under extreme energy limitation□: a synthesis of laboratory- and field-based investigations 1–41.

45. Hoshino T, Toki T, Ijiri A, Morono Y, Machiyama H, Ashi J, Okamura K, Inagaki F. 2017. Atribacteria from the subseafloor sedimentary biosphere disperse to the hydrosphere through submarine mud volcanoes. Front Microbiol 8:1–14.

46. Kopf SH, Sessions AL, Cowley ES, Reyes C, Sambeek L Van, Hu Y, Orphan VJ, Kato R, Newman DK. 2015. Trace incorporation of heavy water reveals slow and heterogeneous pathogen growth rates in cystic fibrosis sputum. Proc Natl Acad Sci USA 113:E110–E116.

47. Brock TD. 1967. Bacterial growth rate in the sea: direct analysis by thymidine autoradiography. Science 155:81–83.

48. Imachi H, Nobu MK, Nakahara N, Morono Y, Ogawara M, Takaki Y, Takano Y, Uematsu K, Ikuta T, Ito M, Matsui Y, Miyazaki M, Murata K, Saito Y, Sakai S, Song C, Tasumi E, Yamanaka Y, Yamaguchi T, Kamagata Y, Tamaki H, Takai K. 2019. Isolation of an archaeon at the prokaryote-eukaryote interface. bioRxiv http://dx.

49. Katayama T, Nobu MK, Kusada H, Meng X-Y, Yoshioka H, Kamagata Y, Tamaki H. 2019. Membrane-bounded nucleoid discovered in a cultivated bacterium of the candidate phylum “Atribacteria.” bioRxiv http://dx.

50. Morono Y, Terada T, Nishizawa M, Ito M, Hillion F, Takahata N, Sano Y, Inagaki F. 2011. Carbon and nitrogen assimilation in deep subseafloor microbial cells. Proc Natl Acad Sci U S A 108:18295–300.

51. Hatzenpichler R, Scheller S, Tavormina PL, Babin BM, Tirrell DA, Orphan VJ. 2014. In situ visualization of newly synthesized proteins in environmental microbes using amino acid tagging and click chemistry. Environ Microbiol 16:2568–2590.

