## Supplementary material for "Evidence for a growth zone for deep subsurface microbial clades in near-surface anoxic sediments": All supplemental information

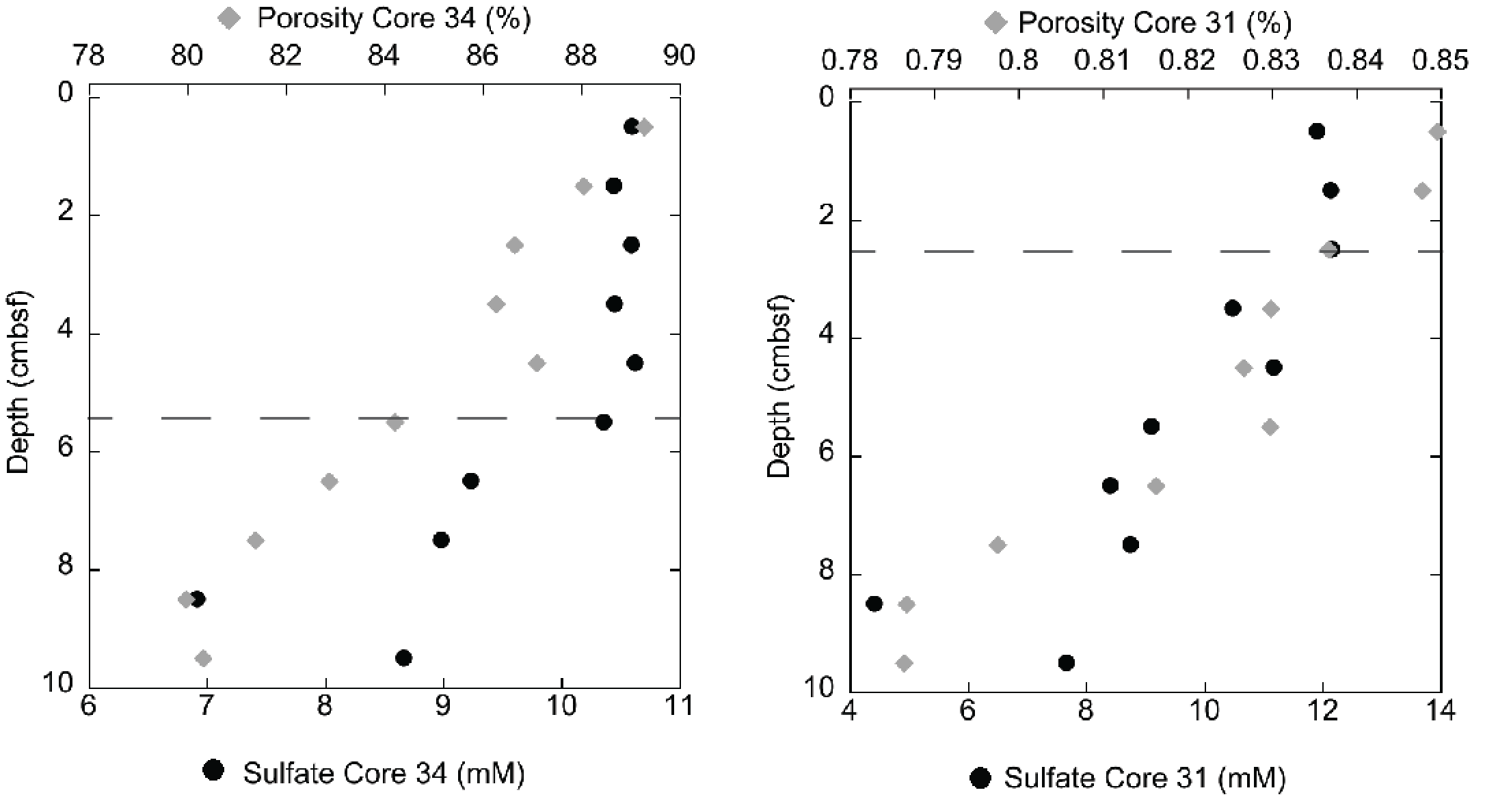

A

B

**Fig. S1 Sulfate and porosity in White Oak River estuary sediments.** Bioirrigation depth (dashed line) determined where porosity and sulfate first start to decrease consistently with depth in A) core 34, which was adjacent to core 30, and B) core 31 which was adjacent to core 32

Fig. S2. Cell abundance over the top 20 cm of White Oak River estuary sediments from cores taken in 2012 and 2013 (used for all analyses in this study except for Table 2).

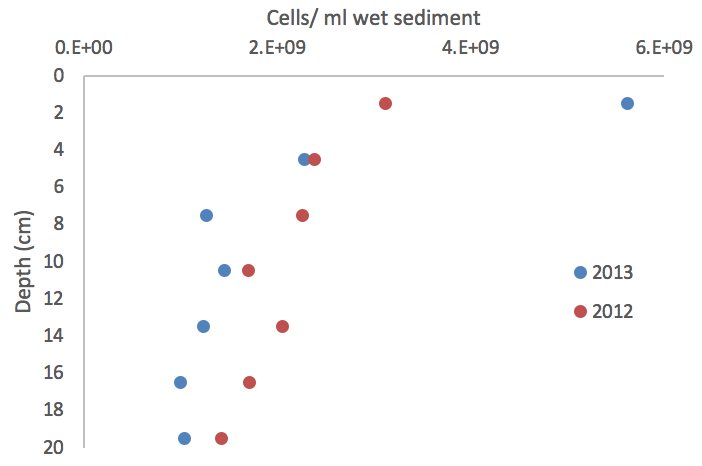

**Figure S3. Archaea, Bathyarchaeota, and MBG-D, but not bacteria, increase in the 3 cm below the depth of bioirrigation in White Oak River estuary sediments** in A-B) core 30 and C-E) core 32, using the product of the fraction of 16S rRNA gene read abundance and cell counts (FRAxC) and quantitative PCR (qPCR) measurements. Grey dotted line is the depth of bioirrigation determined in Fig. 1. Note variations in scale in the x axes.

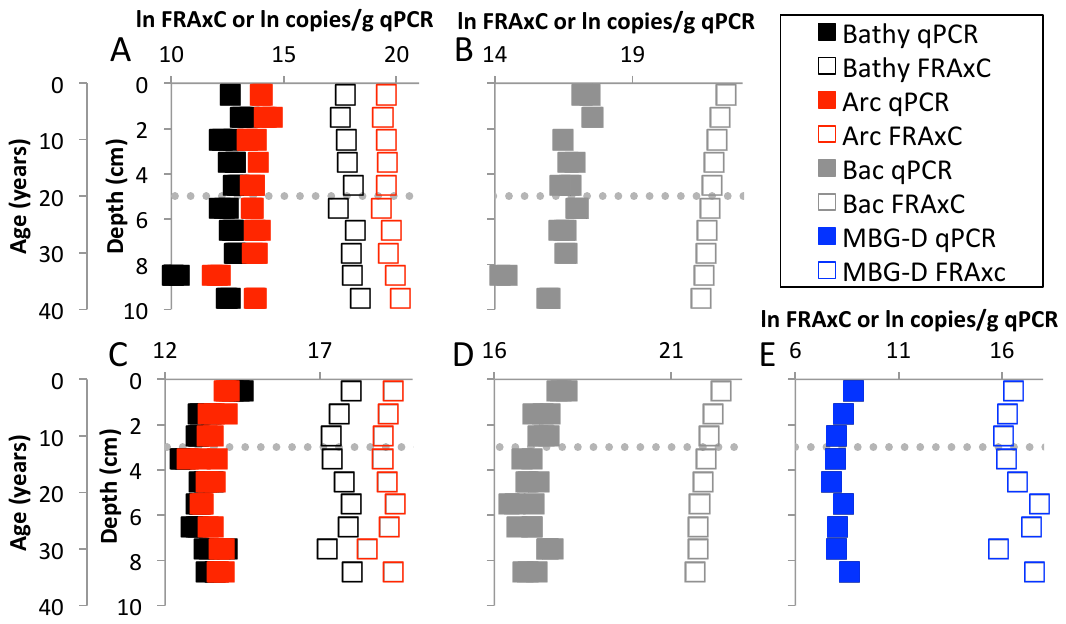

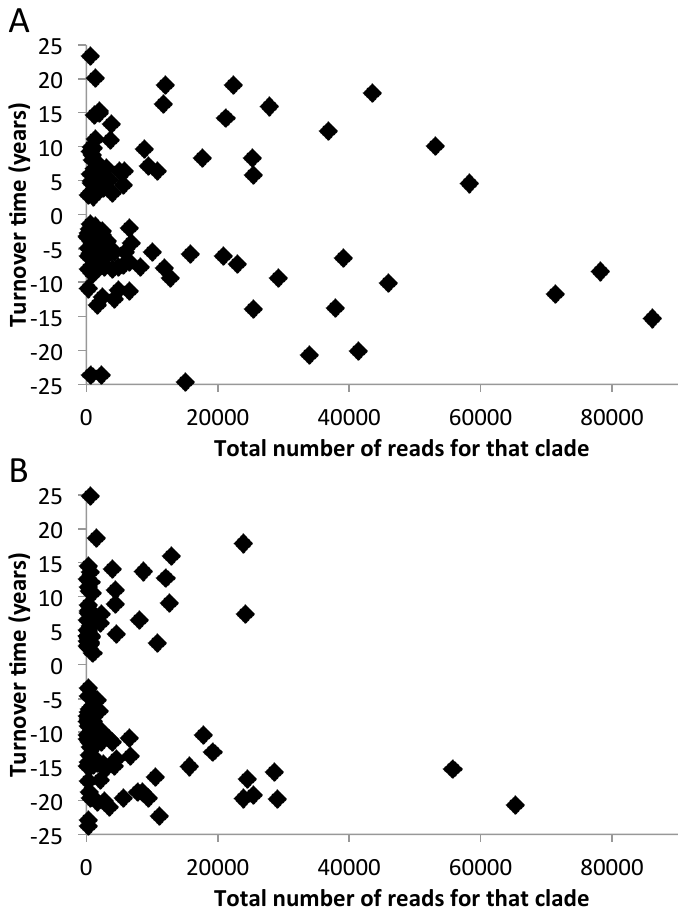

**Figure S4.** **No large trend between turnover time and the number of reads of 16S rRNA genes** for (A) core 30, and (B) core 32. Turnover times calculated from changes in FRAxC with depth and Equation 1.

**Table S1.** Turnover times for each family level clade across the three centimeters below bioirrigation (5.5-7.5 cm for core 30 and 3.5-5.5 cm). These are calculated from FRAxC in White Oak River cores. Also included are the total reads for each clade in each core. Clades are listed in the order of read abundance in core 30.

| **Clades, at the family level** | **Core 30 total reads** | **30.turnover.time 5.5-7.5cm** | **Core 32 total reads** | **32.turnover.time 3.5-5.5cm** |
| --- | --- | --- | --- | --- |
| Gammaproteobacteria_unclassified | 114449 | -17.9 | 70093 | -20.8 |
| JTB255_marine_benthic_group | 86143 | -15.3 | 55688 | -11.3 |
| OM190_unclassified | 78279 | -8.4 | 17770 | -8.3 |
| Planctomycetaceae | 71421 | -11.7 | 65162 | -13.8 |
| Marine_Benthic_Group_D_and_DHVEG-1 | 58184 | 4.6 | 10768 | 3.4 |
| Miscellaneous_Crenarchaeotic_Group | 53029 | 10.1 | 24102 | -12.1 |
| Proteobacteria_unclassified | 45834 | -10.1 | 23832 | 8.9 |
| MSBL9_unclassified | 43459 | 17.9 | 12583 | -13.4 |
| Sandaracinaceae | 41341 | -20.0 | 25302 | -22.1 |
| Halieaceae | 39106 | -6.4 | 29112 | 11.6 |
| OM1_clade | 37800 | -13.7 | 28508 | -13.2 |
| Syntrophobacteraceae | 36722 | 12.3 | 23890 | -13.5 |
| PAUC43f_marine_benthic_group_unclassified | 33864 | -20.6 | 19256 | -11.5 |
| Subgroup_6_unclassified | 29176 | -9.3 | 15665 | -9.9 |
| Desulfarculaceae | 27821 | 15.9 | 12015 | -11.1 |
| Dehalococcoidia_unclassified | 25424 | 5.8 | 4521 | 18.4 |
| Phycisphaeraceae | 25365 | -13.9 | 6530 | 5.0 |
| Marine_Benthic_Group_B_unclassified | 25245 | 8.3 | 8007 | -8.6 |
| Woesearchaeota_(DHVEG-6)_unclassified | 22926 | -7.3 | 9501 | 7.8 |
| Sva0485_unclassified | 22361 | 19.0 | 12871 | -13.4 |
| CCA47 | 21202 | 14.2 | 4423 | 11.3 |
| BD7-8_marine_group_unclassified | 20792 | -6.1 | 10997 | -14.6 |
| Aminicenantes_unclassified | 17619 | 8.3 | 8636 | -13.0 |
| Rhodospirillaceae | 15761 | -5.9 | 7834 | -14.0 |
| Kazan-1B-37_unclassified | 14961 | -24.6 | 3491 | -12.5 |
| OPB35_soil_group_unclassified | 12744 | -9.3 | 10457 | -16.4 |
| TA06_unclassified | 11995 | 19.2 | 3960 | -11.9 |
| SAR324_clade(Marine_group_B)_unclassified | 11852 | -7.9 | 8398 | 21.1 |
| Group_C3_unclassified | 11657 | 16.3 | 4428 | -13.0 |
| vadinBA26_unclassified | 10671 | 6.5 | 2106 | 14.9 |
| Chromatiaceae | 9983 | -5.5 | 5583 | 7.1 |
| GIF3_unclassified | 9434 | 7.2 | 2304 | -13.4 |
| Phycisphaerales_unclassified | 8836 | 9.7 | 1064 | -17.8 |
| Myxococcales_unclassified | 8233 | -7.7 | 3443 | 9.1 |
| Rhodobacteraceae | 6853 | -4.1 | 4538 | 8.3 |
| MG_1_Unknown_Family | 6574 | -1.9 | 6629 | -8.7 |
| Haliangiaceae | 6492 | -7.0 | 2655 | -9.6 |
| Nitrosomonadaceae | 6482 | -11.1 | 1583 | -10.4 |
| BD7-11_unclassified | 5781 | -5.4 | 1927 | -10.2 |
| Candidate_division_OP3_unclassified | 5759 | 6.4 | 1213 | -10.8 |
| Kazan-3B-09_unclassified | 5604 | 4.4 | 975 | -13.7 |
| Gemmatimonadaceae | 5599 | -7.4 | 2182 | -8.5 |
| MSBL5_unclassified | 5009 | 6.4 | 726 | -5.9 |
| Rhodothermaceae | 4846 | -7.6 | 3940 | 7.5 |
| MidBa8 | 4746 | -11.1 | 2683 | -17.8 |
| PAUC34f_unclassified | 4318 | -12.4 | 1664 | 8.3 |
| Subgroup_26_unclassified | 4235 | -5.5 | 2075 | -8.9 |
| Bdellovibrionaceae | 3867 | -8.0 | 1416 | -21.8 |
| AMOS1A-4113-D04 | 3850 | 3.2 | 537 | 4.5 |
| Aerophobetes_unclassified | 3806 | 13.4 | 907 | -8.9 |
| MSB-4B10 | 3684 | 11.1 | 347 | -13.6 |
| Saprospiraceae | 3626 | -5.2 | 4292 | -21.9 |
| FW22_unclassified | 3478 | 5.1 | 513 | -8.9 |
| Oceanospirillaceae | 3109 | -3.9 | 1593 | -12.1 |
| KD3-62_unclassified | 3015 | 6.9 | 578 | -17.5 |
| Verrucomicrobiaceae | 2778 | -4.3 | 2135 | -7.6 |
| Nannocystaceae | 2772 | -6.2 | 1015 | 3.5 |
| S0134_terrestrial_group_unclassified | 2718 | -7.7 | 972 | 7.5 |
| Rhizobiales_unclassified | 2571 | -7.5 | 2830 | 9.3 |
| VHS-B3-70 | 2520 | 4.0 | 495 | -11.0 |
| GR-WP33-58 | 2428 | -2.5 | 1274 | 3.7 |
| Spongiibacteraceae | 2407 | -5.7 | 1149 | -10.8 |
| Cytophagales_unclassified | 2391 | -2.9 | 645 | 3.4 |
| UASB-TL25 | 2364 | -12.1 | 1220 | -7.8 |
| Synergistaceae | 2353 | -23.7 | 584 | -7.0 |
| Cellvibrionales_unclassified | 2216 | -3.3 | 135 | -6.4 |
| Eel-36e1D6 | 2065 | -4.9 | 480 | -17.9 |
| Bacteroidetes_vadinHA17_unclassified | 2029 | 15.3 | 1434 | -11.2 |
| GIF9_unclassified | 2022 | 5.5 | 525 | 6.5 |
| LCP-89_unclassified | 1870 | 15.0 | 413 | -14.7 |
| 20a-9_unclassified | 1818 | 7.1 | 257 | -8.7 |
| Bacteroidia_unclassified | 1705 | 5.3 | 24501 | -10.7 |
| TK10_unclassified | 1703 | -5.5 | 1109 | -10.6 |
| Subgroup_10_unclassified | 1650 | -13.3 | 482 | -9.4 |
| KF-JG30-B3 | 1553 | -2.7 | 1662 | -10.6 |
| MSB-5B2_unclassified | 1534 | 4.2 | 753 |  |
| OM182_clade | 1376 | -1.7 | 1226 | -11.4 |
| VHS-B4-70 | 1358 | -7.3 | 547 | -10.1 |
| Marine_Group_III | 1348 | 3.7 | 246 | -22.4 |
| Thaumarchaeota_unclassified | 1320 | 7.9 | 334 | 4.9 |
| Napoli-4B-65_unclassified | 1306 | 5.6 | 160 | 14.5 |
| SB-5_unclassified | 1261 | 11.1 | 818 | 3.7 |
| Fe-A-9_unclassified | 1242 | 4.5 | 214 | -11.0 |
| ANT06-05 | 1216 | 20.2 | 555 | -7.3 |
| Euryarchaeota_unclassified | 1215 | 4.4 | 217 | -4.6 |
| PeM15_unclassified | 1160 | -3.4 | 775 | 4.8 |
| ODP1230B30.09_unclassified | 1139 | 6.8 | 155 | -9.2 |
| SHA-43_unclassified | 1120 | 14.8 | 347 | -8.4 |
| JL-ETNP-Z39_unclassified | 1101 | 9.8 | 448 | 2.8 |
| B1-7BS_unclassified | 1066 | 2.7 | 923 | 11.1 |
| Opitutaceae | 964 | -8.6 | 747 | 3.8 |
| S085_unclassified | 941 | 8.1 | 295 | -17.9 |
| Acidobacteria_subgroup4_Unknown_Family | 939 | -4.8 | 804 | 14.0 |
| T9d | 905 | -2.0 | 865 | 4.7 |
| ODP1230B30.02_sediment_group | 879 | 8.8 | 136 | 5.5 |
| SPOTSOCT00m83_unclassified | 871 | -4.0 | 544 | -4.0 |
| Sh765B-AG-111_unclassified | 800 | 5.0 | 108 | 4.7 |
| Oligoflexaceae | 785 | -2.3 | 488 | 9.7 |
| SC-I-84_unclassified | 780 | -3.9 | 375 | 9.9 |
| OPB41_unclassified | 772 | 9.9 | 619 | 1.9 |
| Miscellaneous_Euryarchaeotic_Group(MEG)_unclassified | 760 | 4.7 | 199 | -5.4 |
| Cryomorphaceae | 758 | -2.1 | 225 | 15.7 |
| MSB-1E8 | 686 | -5.6 | 265 | -8.4 |
| B276-D12_unclassified | 626 | -23.7 | 227 | -10.2 |
| Thermoflexaceae | 616 | 6.0 | 194 | -13.4 |
| JTB23_unclassified | 592 | -1.4 | 488 | -6.4 |
| VC2.1_Arc6 | 572 | 23.4 | 117 | -8.2 |
| Sphingobacteriales_unclassified | 571 | -2.6 | 665 | 17.2 |
| possible_order_07_unclassified | 570 | 9.3 | 156 | 3.1 |
| Hyd24-12_unclassified | 536 | 3.2 | 102 | -7.7 |
| 0319-6A21 | 529 | -5.3 | 161 | -17.9 |
| PL-11B10 | 524 | -5.9 | 212 | -14.8 |
| Subgroup_15_unclassified | 497 | -2.8 | 433 | -6.8 |
| Acetobacteraceae | 459 | -2.1 | 510 | 4.3 |
| Erythrobacteraceae | 448 | -3.4 | 434 | -12.8 |
| Puniceicoccaceae | 412 | -2.8 | 196 | -13.0 |
| Sphingomonadales_unclassified | 370 | -3.1 | 362 | -8.9 |
| Granulosicoccaceae | 334 | -8.0 | 344 | -11.0 |
| Rhodocyclaceae | 289 | -6.2 | 457 | 7.8 |
| Rikenellaceae | 283 | 2.9 | 295 | -10.2 |
| Opitutae_unclassified | 251 | -4.9 | 122 | -8.3 |
| Acidobacteria_subgroup3_Unknown_Family | 240 | -10.9 | 242 | -7.5 |
| SC3-20_unclassified | 194 | -2.7 | 221 | -5.7 |
| Piscirickettsiaceae | 176 | -3.4 | 158 | -10.8 |
| Verrucomicrobia_unclassified | 174 | -3.1 | 140 | -4.1 |

**Table S2.** Turnover times calculated with FRAxC for each family level clade using timepoints between 40 and 802 days (40, 47, 54, 61, 75, 80, 86, 94, 107, 114, 122, and 802 days). Clades are organized by their abundance, shown by ln(FRAxC).

| **Clades, at the family level** | **Incu-bation 2 ln(FRAxC)** | **Incu-bation 2 doubling.time** | **Incu-bation 3 ln(FRAxC)** | **Incu-bation 3 doubling.time** |
| --- | --- | --- | --- | --- |
| Deinococcaceae | 19.8 | -0.5 | 20.0 | -0.6 |
| GIF3_unclassified | 19.1 | -1.3 | 19.1 | -1.7 |
| Shewanellaceae | 18.7 | -0.5 | 19.4 | -0.4 |
| FW113_unclassified | 18.7 | -0.4 | 18.9 | -0.5 |
| Desulfomicrobiaceae | 18.4 | -0.6 | 18.7 | -0.6 |
| Synergistaceae | 18.3 | -1.3 | 18.7 | -1.3 |
| CK-1C4-49_unclassified | 18.3 | -0.4 | 18.2 | -0.4 |
| Marine_Benthic_Group_D_and_DHVEG-1 | 18.3 | 3.4 | 18.2 | 4.4 |
| Epsilonproteobacteria_unclassified | 18.1 | -0.6 | 18.4 | -0.6 |
| Burkholderiales_unclassified | 18.0 | -0.4 | 18.5 | -0.4 |
| Oligosphaeraceae | 18.0 | -0.8 | 18.0 | -1.0 |
| Archaea_unclassified | 18.0 | -1.1 | 18.4 | -1.1 |
| Woesearchaeota_(DHVEG-6)_unclassified | 17.9 | -0.7 | 18.4 | -0.7 |
| Subgroup_2_unclassified | 17.8 | -0.7 | 18.0 | -0.7 |
| Psychromonadaceae | 17.8 | -0.8 | 18.6 | -0.6 |
| BD7-8_marine_group_unclassified | 17.7 | -0.5 | 18.2 | -0.4 |
| GIF9_unclassified | 17.6 | -0.6 | 18.4 | -0.5 |
| Syntrophomonadaceae | 17.5 | -0.6 | 17.9 | -0.7 |
| Rhodospirillales_unclassified | 17.5 | -0.6 | 17.8 | -0.6 |
| Methylococcales_unclassified | 17.4 | -0.4 | 17.7 | -0.7 |
| SAR324_clade(Marine_group_B)_unclassified | 17.4 | -0.4 | 17.9 | -0.4 |
| Flavobacteriales_unclassified | 17.3 | -0.4 | 17.6 | -0.5 |
| Napoli-4B-65_unclassified | 17.3 | -0.7 | 17.9 | -0.6 |
| Order_III_unclassified | 17.2 | -0.3 | 17.5 | -0.3 |
| Marine_Benthic_Group_B_unclassified | 17.1 | -1.6 | 16.9 | -1.0 |
| CCA47 | 17.1 | -2.4 | 17.1 | -3.0 |
| 10bav-F6_unclassified | 17.1 | -0.4 | 17.3 | -0.4 |
| TK34 | 17.0 | -0.5 | 17.0 | -0.6 |
| Legionellaceae | 17.0 | -0.4 | 16.9 | -0.3 |
| Cytophagales_unclassified | 17.0 | -0.4 | 17.4 | -0.4 |
| JG34-KF-361 | 17.0 | -0.5 | 17.3 | -0.4 |
| SB1-18_unclassified | 16.9 | -0.5 | 17.6 | -0.5 |
| Bdellovibrionaceae | 16.9 | -0.3 | 17.8 | -0.3 |
| Family_XII | 16.9 | -0.4 | 16.8 | -0.4 |
| Subgroup_5_unclassified | 16.8 | -0.4 | 17.1 | -0.4 |
| CS-B046_unclassified | 16.8 | -2.0 | 16.9 | -2.1 |
| Dictyoglomaceae | 16.8 | -0.8 | 16.9 | -0.8 |
| OPB41_unclassified | 16.8 | -0.6 | 16.9 | -0.7 |
| Geobacteraceae | 16.8 | -0.5 | 17.3 | -0.5 |
| EC3_unclassified | 16.7 | -0.4 | 16.8 | -0.3 |
| JL-ETNP-Z39_unclassified | 16.6 | -0.8 | 17.3 | -1.2 |
| Armatimonadetes_unclassified | 16.6 | -0.7 | 16.6 | -0.8 |
| TK85 | 16.6 | -0.4 | 17.0 | -0.4 |
| Nitrosomonadaceae | 16.6 | -0.3 | 16.8 | -0.3 |
| Pseudoalteromonadaceae | 16.6 | -1.6 | 16.7 | -1.9 |
| Rhodothermaceae | 16.5 | -0.5 | 16.4 | -0.7 |
| Haloplasmataceae | 16.4 | -1.0 | 16.8 | -1.2 |
| Halieaceae | 16.4 | -0.4 | 16.5 | -0.4 |
| TPD-58_unclassified | 16.4 | -0.6 | 17.1 | -0.6 |
| Dehalococcoidia_unclassified | 16.3 | -0.7 | 16.8 | -0.6 |
| ODP1230B30.09_unclassified | 16.3 | -0.5 | 16.5 | -0.5 |
| NS9_marine_group | 16.3 | -0.4 | 16.4 | -0.5 |
| Eubacteriaceae | 16.3 | -0.5 | 16.7 | -0.5 |
| c5LKS8_unclassified | 16.2 | -0.6 | 16.6 | -0.7 |
| Helicobacteraceae | 16.2 | -0.3 | 16.7 | -0.3 |
| Schleiferiaceae | 16.2 | -0.3 | 16.4 | -0.4 |
| KD3-62_unclassified | 16.2 | -0.4 | 16.0 | -0.4 |
| Cryomorphaceae | 16.2 | -0.4 | 16.3 | -0.4 |
| Cyanobacteria_unclassified | 16.2 | -1.4 | 16.8 | -1.0 |
| Nannocystaceae | 16.1 | -0.5 | 16.5 | -0.5 |
| Fibrobacterales_unclassified | 16.1 | -0.4 | 16.1 | -0.3 |
| ST-12K33 | 16.1 | -0.3 | 15.8 | -0.4 |
| 0319-6A21 | 16.0 | -0.6 | 16.3 | -0.6 |
| Holosporaceae | 16.0 | 3.7 | 16.6 | 3.7 |
| Piscirickettsiaceae | 16.0 | -0.5 | 16.1 | -0.7 |
| C86_unclassified | 16.0 | -0.9 | 15.8 | -1.5 |
| EF100-94H03 | 16.0 | -0.5 | 16.5 | -0.6 |
| Unknown_Family | 16.0 | -0.4 | 16.3 | -0.4 |
| Spirochaetales_unclassified | 16.0 | -0.7 | 16.0 | -0.7 |
| BIrii41 | 15.9 | -0.6 | 16.5 | -0.7 |
| Miscellaneous_Crenarchaeotic_Group_unclassified | 15.9 | 6.4 | 15.5 | 6.3 |
| Rs-M47_unclassified | 15.9 | -3.7 | 16.0 | -2.1 |
| Subgroup_25_unclassified | 15.9 | -0.7 | 16.4 | -0.7 |
| Sphingobacteriales_unclassified | 15.9 | -0.9 | 16.0 | -2.2 |
| Nitrospiraceae | 15.8 | -0.4 | 16.8 | -0.4 |
| OM182_clade | 15.8 | -0.4 | 16.1 | -0.5 |
| ARKDMS-49_unclassified | 15.8 | -0.6 | 16.0 | -0.7 |
| Oceanospirillaceae | 15.8 | -0.3 | 16.0 | -0.3 |
| Verrucomicrobiaceae | 15.7 | -0.3 | 16.0 | -0.3 |
| MSBL5_unclassified | 15.7 | -0.4 | 15.9 | -0.3 |
| GR-WP33-30_unclassified | 15.7 | -0.3 | 15.4 | -0.4 |
| SBYZ-984 | 15.6 | -0.4 | 16.2 | -0.5 |
| Gemmatimonadaceae | 15.6 | -0.4 | 16.4 | -0.4 |
| LWSR-14 | 15.6 | -0.3 | 16.4 | -0.3 |
| Lentisphaerae_RFP12_gut_group_unclassified | 15.6 | -1.3 | 15.4 | -3.2 |
| LD1-PA38_unclassified | 15.6 | -2.5 | 15.2 | 1.7 |
| AKIW1012 | 15.6 | -1.0 | 15.5 | -1.8 |
| Trueperaceae | 15.6 | -0.2 | 15.8 | -0.3 |
| Planctomycetacia_unclassified | 15.6 | -1.7 | 15.7 | -2.6 |
| Unknown_Family | 15.5 | -0.6 | 16.0 | -0.5 |
| DUNssu371 | 15.5 | -0.5 | 15.6 | -0.5 |
| Lentisphaerae_unclassified | 15.5 | -1.3 | 16.1 | -1.2 |
| Cyclobacteriaceae | 15.5 | -0.3 | 16.0 | -0.3 |
| Brocadiaceae | 15.5 | -0.9 | 15.6 | -0.7 |
| Xanthobacteraceae | 15.5 | -0.3 | 15.6 | -0.4 |
| Rubrobacteriaceae | 15.5 | -0.5 | 15.9 | -0.4 |
| Group_C3_unclassified | 15.5 | 2.1 | 15.3 | 1.8 |
| AT-s3-44 | 15.5 | -0.4 | 15.9 | -0.4 |
| BD2-7 | 15.4 | -1.9 | 15.7 | -2.5 |
| Campylobacterales_unclassified | 15.4 | -0.4 | 15.9 | -0.3 |
| Puniceicoccaceae | 15.4 | -0.3 | 16.2 | -0.3 |
| SHA-43_unclassified | 15.4 | -1.5 | 15.7 | -1.8 |
| Solirubrobacteraceae | 15.4 | -0.4 | 16.0 | -0.4 |
| Thermoplasmatales_unclassified | 15.4 | -1.8 | 15.5 | -1.7 |
| Acidothermaceae | 15.4 | -0.6 | 15.9 | -0.6 |
| Rhodobiaceae | 15.4 | -0.4 | 15.3 | -0.4 |
| Syntrophobacteraceae | 15.3 | -0.3 | 15.3 | -0.3 |
| Bradyrhizobiaceae | 15.3 | -0.5 | 15.2 | -0.6 |
| Idiomarinaceae | 15.3 | -0.4 | 15.7 | -0.4 |
| Kazan-3A-21_unclassified | 15.3 | 1.1 | 14.9 | 0.9 |
| Aerophobetes_unclassified | 15.2 | -0.5 | 15.2 | -0.6 |
| OPB35_soil_group_unclassified | 15.2 | -0.4 | 15.3 | -0.3 |
| 20c-4 | 15.2 | 2.4 | 14.7 | 1.7 |
| Porphyromonadaceae | 15.2 | -0.3 | 15.3 | -0.5 |
| HOC36_unclassified | 15.2 | -0.5 | 15.3 | -0.6 |
| Desulfuromonadaceae | 15.1 | -0.3 | 16.2 | 0.1 |
| Marine_Hydrothermal_Vent_Group(MHVG)_unclassified | 15.1 | -1.5 | 14.9 | -2.9 |
| Subgroup_9_unclassified | 15.1 | -0.6 | 15.2 | -0.6 |
| Parcubacteria_unclassified | 15.1 | -1.3 | 15.6 | -1.5 |
| T9d | 15.1 | -1.2 | 15.1 | -1.4 |
| Methanosaetaceae | 15.1 | 0.8 | 15.0 | 0.4 |
| ODP1230B30.02_sediment_group | 15.1 | -0.7 | 15.3 | -0.8 |
| Coriobacteriaceae | 15.0 | -0.2 | 15.1 | -0.3 |
| Fusobacteriaceae | 15.0 | -0.8 | 15.9 | -0.7 |
| Lineage_IV_unclassified | 15.0 | -1.9 | 15.2 | -1.6 |
| Deferribacteraceae | 15.0 | -0.5 | 15.4 | -0.5 |
| Methanomicrobiales_unclassified | 15.0 | 0.7 | 15.9 | 0.6 |
| MSBL2_unclassified | 15.0 | -0.9 | 15.6 | -0.7 |
| Obscuribacterales_unclassified | 14.9 | -0.6 | 15.9 | -0.5 |
| MSBL3_unclassified | 14.9 | -1.8 | 15.5 | -1.8 |
| Mollicutes_unclassified | 14.8 | -0.6 | 15.4 | -0.7 |
| LH041 | 14.8 | -1.2 | 14.5 | -3.2 |
| Haliangiaceae | 14.8 | -0.4 | 15.3 | -0.5 |
| MSBL9_unclassified | 14.8 | -3.2 | 14.6 | -2.9 |
| LCP-89_unclassified | 14.8 | -0.8 | 15.3 | -0.7 |
| ANME-1b | 14.8 | 0.4 | 14.8 | 0.4 |
| B103G10_unclassified | 14.7 | -1.6 | 14.3 | -3.9 |
| Unknown_Family | 14.7 | -0.4 | 14.7 | -0.4 |
| Ktedonobacteria_unclassified | 14.7 | -0.5 | 15.1 | -0.4 |
| Acetobacteraceae | 14.7 | -0.4 | 15.2 | -0.5 |
| Unknown_Family | 14.7 | -0.4 | 15.3 | -0.4 |
| Ancient_Archaeal_Group(AAG)_unclassified | 14.7 | -2.0 | 14.6 | -1.2 |
| Nitrospinaceae | 14.7 | -0.5 | 14.7 | -0.4 |
| TA18_unclassified | 14.7 | -0.7 | 14.9 | -0.6 |
| TK10_unclassified | 14.7 | -0.4 | 15.0 | -0.3 |
| Marine_Group_III | 14.7 | 1.4 | 14.3 | 1.5 |
| Pla4_lineage_unclassified | 14.6 | -0.7 | 14.9 | -0.9 |
| Thermomicrobia_unclassified | 14.6 | -0.3 | 14.8 | 0.1 |
| SPOTSOCT00m83_unclassified | 14.6 | -0.7 | 14.9 | -0.6 |
| IheB3-7 | 14.6 | -0.4 | 14.9 | -0.4 |
| BD7-11_unclassified | 14.6 | -1.5 | 15.3 | -1.1 |
| Hyd24-12_unclassified | 14.6 | -1.1 | 14.7 | -1.2 |
| Bacteroidetes_vadinHA17_unclassified | 14.6 | -0.3 | 14.3 | -0.4 |
| Burkholderiaceae | 14.6 | -0.3 | 14.9 | -0.4 |
| Moritellaceae | 14.5 | -0.4 | 14.7 | -0.4 |
| Nocardiaceae | 14.5 | -0.3 | 14.2 | 0.1 |
| ML-A-10_unclassified | 14.5 | -2.3 | 14.8 | -2.6 |
| VC2.1_Arc6 | 14.5 | -1.7 | 14.5 | -0.8 |
| Oligoflexaceae | 14.5 | -0.4 | 14.6 | -0.4 |
| Chthoniobacterales_unclassified | 14.5 | -0.4 | 14.8 | -0.5 |
| SHA-109_unclassified | 14.5 | -0.4 | 14.5 | -0.6 |
| vadinHA49_unclassified | 14.5 | -1.3 | 14.5 | -1.8 |
| Subgroup_7_unclassified | 14.5 | -0.4 | 14.9 | -0.3 |
| Acetothermia_unclassified | 14.4 | -6.0 | 14.4 | -4.3 |
| Oceanospirillales_unclassified | 14.4 | -0.6 | 14.9 | -0.8 |
| Omnitrophica_unclassified | 14.4 | -0.6 | 15.1 | -0.7 |
| Brevinemataceae | 14.3 | -0.3 | 14.7 | 0.3 |
| MSB-1E8 | 14.3 | -0.3 | 14.5 | -0.3 |
| Phyllobacteriaceae | 14.3 | -0.8 | 14.7 | -1.4 |
| Desulfobacterales_unclassified | 14.3 | -0.3 | 14.7 | -1.8 |
| MgMjR-022 | 14.3 | -0.5 | 14.4 | -0.5 |
| Alcanivoracaceae | 14.3 | -0.3 | 14.6 | -0.3 |
| Aminicenantes_unclassified | 14.3 | -0.5 | 15.4 | -0.6 |
| Proteobacteria_unclassified | 14.2 | -0.9 | 14.3 | -2.2 |
| B2706-C7 | 14.2 | -0.6 | 14.7 | -0.6 |
| S-BQ2-57_soil_group_unclassified | 14.2 | -0.4 | 14.4 | -0.5 |
| 64K2 | 14.2 | -1.1 | 14.6 | -0.5 |
| Corynebacteriaceae | 14.2 | -0.3 | 14.1 | 0.1 |
| Rickettsiales_unclassified | 14.2 | -0.3 | 14.3 | -0.7 |
| Rikenellaceae | 14.1 | -0.3 | 14.6 | -0.4 |
| 09D2Z46 | 14.1 | -2.6 | 14.2 | -3.5 |
| Sva0071_unclassified | 14.1 | -0.5 | 14.3 | -0.6 |
| P._palm_C-A_51 | 14.1 | -0.5 | 14.3 | -0.3 |
| Euryarchaeota_unclassified | 14.1 | -1.2 | 14.6 | -1.3 |
| 20a-9_unclassified | 14.1 | 0.8 | 14.2 | 0.7 |
| Methylobacteriaceae | 14.1 | -0.7 | 14.1 | -0.7 |
| Cyanobacteria_unclassified | 14.1 | -0.5 | 14.5 | -0.8 |
| Saccharibacteria_unclassified | 14.0 | -0.6 | 14.2 | -0.8 |
